# Comparative analysis of STP6 and STP10 unravels molecular selectivity in Sugar Transport Proteins

**DOI:** 10.1101/2024.07.19.604285

**Authors:** Camilla Gottlieb Andersen, Laust Bavnhøj, Søren Brag, Anastasiia Bohush, Adriana Chrenková, Jan Heiner Driller, Bjørn Panyella Pedersen

**Author notes:** To whom correspondence should be addressed. (BPP).

## Abstract

The distribution of sugars is crucial for plant energy, signaling, and defense mechanisms. Sugar Transport Proteins (STPs) are Sugar Porters that mediate proton-driven cellular uptake of glucose. Some STPs also transport fructose, while others remain highly selective for only glucose. What determines this selectivity, allowing STPs to distinguish between compounds with highly similar chemical composition, remains unknown. Here, we present the structure of *Arabidopsis thaliana* STP6 in an inward occluded conformational state with glucose bound and demonstrate its role as both a glucose and fructose transporter. We perform a comparative analysis of STP6 with the glucose-selective STP10 using *in-vivo* and *in-vitro* systems, demonstrating how different experimental setups strongly influence kinetic transport properties. We analyze the properties of the monosaccharide binding site and show that the position of a single methyl group in the binding site is sufficient to shuffle glucose and fructose specificity, providing detailed insights into the fine-tuned dynamics of affinity-induced specificity for sugar uptake. Altogether these findings enhance our understanding of sugar selectivity in STPs and more broadly Sugar Porter proteins.

**SIGNIFICANCE STATEMENT:** Understanding the mechanisms of sugar transport in plants is essential for advancing agricultural practices and enhancing plant resilience. This study reveals the structural basis of sugar selectivity in Sugar Transport Proteins of Arabidopsis thaliana. By comparing the dual-specific STP6, transporting both glucose and fructose with the glucose-selective STP10 across multiple experimental setups, we show that difference as subtle as the position of a single methyl group in the binding site can control sugar specificity. These findings enhance our understanding of sugar selectivity by Sugar Transport Proteins and more broadly Sugar Porter proteins and lay the groundwork for engineering crops with improved energy efficiency and pathogen resistance.

## INTRODUCTION

Sugar serves both as an energy source and essential signaling molecules controlling tissue development in plants. Cellular uptake of monosaccharides, in particular glucose, is mediated by Sugar Transport Proteins (STPs) that import monosaccharides by proton driven symport, utilizing the proton motive force (PMF) (1–4). Their function is central to correct tissue and organ development. STPs form a distinct plant-specific subgroup in the Sugar Porter (SP) family, which in turn constitutes the largest family within the Major Facilitator Superfamily (MFS), and include human GLUTs (5, 6). 14 STPs are found in *Arabidopsis thaliana* (STP1 to STP14) where they are primarily expressed in symplastically isolated cells in tissues such as seeds, flowers, pollen, and roots (7). They undergo extensive transcriptional regulation during lateral root formation and pollen maturation, essential to ensure correct development (8, 9). In addition, they aid in the plant’s defense machinery against invading pathogens by sequestering monosaccharides from the apoplast and are directly linked to environmental stress responses triggered by challenges such as drought or extreme temperatures (10–16). *A. thaliana* STP10 have provided the foundation for the current understanding of the STP transport mechanism (17, 18). STP10 has been characterized in multiple experimental setups, where it has been shown to transport glucose with high affinity (K_M_^apps^ = 2-20 µM) in a proton dependent manner (17, 19). Two glucose epimers, galactose and mannose, inhibit STP10’s uptake of glucose in competition assays (17). The structures of STP10 solved in complex with glucose, in outward occluded and inward open conformational states, provided detailed insights into the alternating access mechanism of STPs used for transport (17, 18, 20–23). STPs adopt the canonical MFS fold with 12 transmembrane helices (M1-M12) present in two membrane-spanning domains (N and C) (24–26).

The two domains are related by rotational pseudo symmetry, with the sugar binding site sandwiched in-between. A distinct protonation site in M1 (Asp42 in STP10) is linked to the substrate binding site, ensuring tight proton-sugar coupling (17, 18). As in all other SP family proteins, the N and C domains are connected by an intracellular cytosolic helix bundle (ICH) domain (27–30). The ICH domain also holds the SP signature motif, which is often seen to interact with the MFS specific A motif, localized between M2 and M3, M8 and M9 (31). The A motif is involved in forming interdomain salt-bridge networks, known to stabilize the transporter in outward facing conformations (18, 28, 32). Both the A and the SP motifs have been suggested to play a key role in regulating transport kinetics by several studies conducted on SP family members (29, 31, 33). A unique feature of the STPs is the presence of a disulfide bridge present on a small extracellular Lid domain located between M1 and M2, tethering the N domain to the C domain. A previous study has shown that the disulfide bridge is needed for high glucose affinity, possibly helping the Lid domain shield the protonation site from the extracellular environment (17).

Transport activity has been characterized for several STPs across species in different experimental setups. While glucose is the preferred substrate for most STPs, several STPs are promiscuous and transport a range of monosaccharides (4, 7, 34). Notably, the hydrolysis of photosynthesis-derived sucrose in the apoplast yields equimolar amounts of glucose and fructose (35, 36), but only STP6, STP8, and STP13, have been suggested to facilitate significant uptake of fructose, while STP10 remain highly selective for glucose (9, 13, 37).

Here we present a structural and biophysical characterization of *A. thaliana* STP6, including a direct comparative analysis with STP10 addressing differences in substrate specificities (52% sequence identity). With a 3.2 Å crystal structure of STP6 in complex with glucose, we detail a previously unresolved inward occluded state of the STP transport cycle. Our biophysical characterization shows that STP6 and STP10 transports a range of hexoses with varying preference and demonstrates dependency on the PMF. We identify affinity-determining sites and show how the position of a single methyl group can be used to rationally modify the transport affinities of STP6 and STP10 towards glucose and fructose. Finally, our work represents the use of diverse experimental setups in parallel to demonstrate how transporter kinetics are highly dependent on the context of transporter environment, relevant for comparative analysis of membrane transporter biophysics between related targets.

## RESULTS

To enable a strict comparative analysis of STP6 and STP10, we established a shared protocol for expression and purification, to ensure identical baseline working conditions and samples (Supplementary Figure 1). STP6 activity was tested using uptake assays in *Saccharomyces cerevisiae* and compared to previous characterizations of STP10 performed under identical experimental conditions (17, 18). STP6 transports glucose with high affinity, K_M_^app^ = 23.8 µM, in a pH dependent manner (Figure 1A and Supplementary Figure 2), corresponding well with previously reported affinities obtained from yeast uptake assays of STP6 (20 µM) and STP10 (19 µM), as well as the shown pH preference for STP10 (pH 4 – 6.5) (9, 17, 19). STP6-mediated transport is highly dependent on the PMF, as demonstrated by the significant reduction (∼76 % decrease) in the presence of the proton gradient decoupler carbonyl cyanide m-chlorophenyl hydrazone (CCCP) and compromised by a range of inhibitors known to affect SP proteins, including curcumin and phloretin (Supplementary Figure 2) (1–3, 17, 19, 38, 39). Mannose and galactose are able to compete with glucose uptake by STP6 in competition assays and were both shown to be transported with intermediate affinities (K_M_^app^ = 530 µM and K_M_^app^ = 129 µM, respectively) (Figure 1B and Supplementary Figure 2). While xylose and sucrose do not compete with glucose, correlating with the reported competition profile for STP10 and supporting the expected preference for hexoses, competition by fructose is only observed in STP6 (Figure 1B). These data cements STP6 as a proton/hexose symporter, displaying high affinity for glucose with the additional ability to transport fructose.

**Figure 1.**
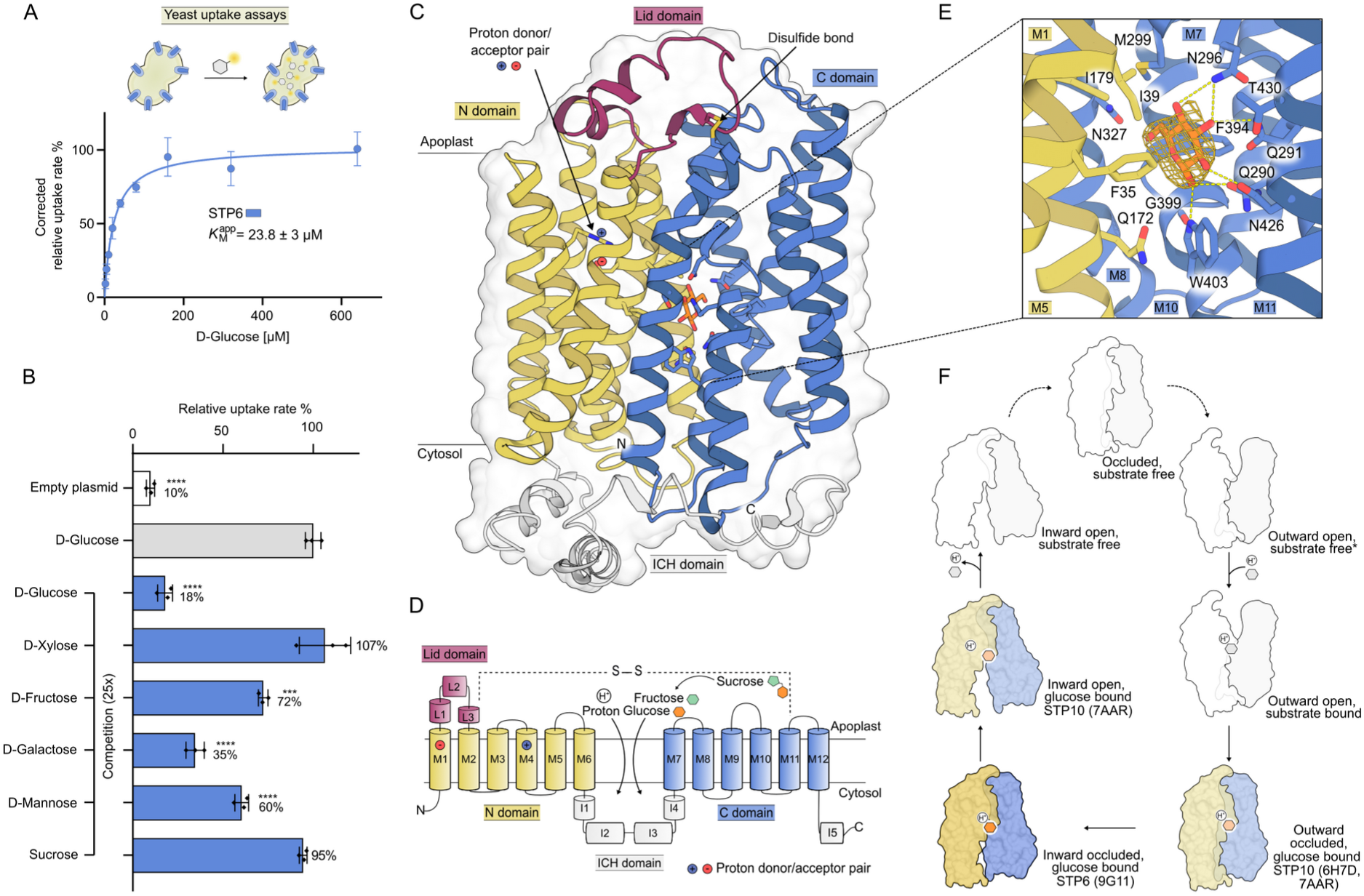
Functional and structural characteristics of STP6. A) Michaelis-Menten analysis of labelled D-glucose uptake by heterologous expression of STP6 in yeast cells. B) Substrate specificity of STP6 assed by competition assay performed in yeast cells with 10 µM glucose and 250 µM of the competing sugar. *P-*values have been obtained by one-way ANOVA analysis ***, *p* = 0.0002 and ****, *p* < 0.0001. C) Cartoon and surface representation of the STP6 crystal structure in an inward occluded conformational state with glucose bound in-between the N domain (yellow) and the C domain (blue). Residues involved in substrate interactions are shown as sticks. The proton donor/acceptor pair and the disulfide bond connecting the Lid domain, and the C domain are highlighted with arrows. D) Schematic representation of STP6 topology. The proton donor/acceptor pair is located in M4 and M1, respectively and the disulfide bond is formed between L3 and M11, as indicated. STP6 transports glucose and fructose, derived from apoplastic sucrose E) Close-up view of the glucose binding site. Interacting residues are shown as sticks and yellow dashes represent hydrogen bonding (2.6-3.6 Å distance). The 2Fo-Fc map for glucose is shown as gold mesh (contoured 2σ). F) Illustration of key conformational states in the STP transport cycle. Binding of substrate takes place in the outward open state, substrate transitioning via occluded states and lastly release is permitted in the inward open state. Following substrate release the transport is reset through substrate empty occluded states before the cycle can be repeated. Experimentally determined states are highlighted with colors and * marks the start of the transport cycle.

Since no experimental structural information of STP6 was available for use in structural comparisons with STP10, we crystallized STP6 in complex with glucose and solved the structure to a resolution of 3.2 Å. STP6 crystallizes in a P3_2_21 space group with a single monomer in the asymmetric unit. The final model includes residues 17 to 492 (of 507 residues total), glucose, cholesteryl hemisuccinate (CHS), and was refined to an R-free of 25.2% (Figure 1C, Supplementary Table 1 and Supplementary Figure 3). STP6 displays the MFS fold, with 12 transmembrane helices (M1-M12) divided between an N domain and a C domain (M1-M6 and M7-M12, respectively). Four of the five-bundle intracellular helix domain (ICH1-5) connects the N and C domains. The extracellular helical Lid domain (L1-3) is found between M1 and M2, where it is covalently linked to the C domain by a disulfide bridge formed between Cys72 (L3) and Cys442 (M11) (Figure 1C and D). The positioning of three CHS molecules mimics the inner leaflet of the plasma membrane (Supplementary Figure 3). One of the CHS head groups interacts with the A motif through Arg340 (M8) (Supplementary Figure 3). STP6 adopts a substrate-bound inward occluded conformational state with glucose bound in the central cavity (Figure 1C, E and F). The majority of the interactions between glucose and binding pocket are to the C domain, by a distinct network of mainly polar interactions to Gln290, Gln291, Asn296, Met299, Asn327, Gly399, Trp403, Asn426 and Thr430 (Figure 1E). The N domain binds the glucose molecule predominantly through hydrophobic interactions mediated by Ile39 and Ile179 and T-shaped π-stacking between Phe35 and the glucosyl ring (Figure 1E). In STP6, the protonation site is separated from the binding site, with the proton donor/acceptor, Asp38, positioned on the M1 helix. Arg137 is found in close spatial vicinity to Asp38, sitting on M4, wherefrom it controls the pKa of Asp38. Together, they constitute a classical proton donor-acceptor pair (Figure 1C and D) (40). The 4.3 Å distance between Asp38 and Arg137 suggests protonation of Asp38, as the pair otherwise would be forming a salt bridge (>3 Å) (Supplementary figure 4) (18). In addition, Asp38 is seen to point away from Arg137, which is one of two protonation orientations that has currently been observed for the proton donor-acceptor pair in structures of STPs (Supplementary figure 4) (18). Collectively, these observations reflect the structural arrangement of the key residues needed to ensure tight coupling between glucose-binding and protonation of Asp38, previously described for STP10 and characteristic for proton-coupled symporters in the SP family (17, 18, 29).

Compared to the known structures of STP10 determined in the outward occluded and inward open states, STP6 adopts a distinct inward occluded conformational state (Figure 1F). The inward occluded state shares many of the characteristics observed in the inward open state of STP10, as demonstrated by plotting the domain-paired root mean square deviation (RMSD) between the corresponding C-alpha backbone positions (N domain RMSD(Cα) 1.141 Å, C domain RMSD(Cα) 0.734 Å) (Figure 2A and Supplementary Figure 4). The low RMSD of the C domain reflects that both of the inward states exhibit the same helical rearrangement when progressing from the outward state. In this process the C domain inverts into a “substrate release conformation”, which includes outwards kinking of M11b that in turn pushes the core of M7 down towards the substrate binding site, allowing Met299 and Phe300 to contact M1 and form a hydrophobic outer gate (Figure 2B). This joint movement forces the Lid domain to bend away from the body of the transporter and adopt a more open conformation.

**Figure 2.**
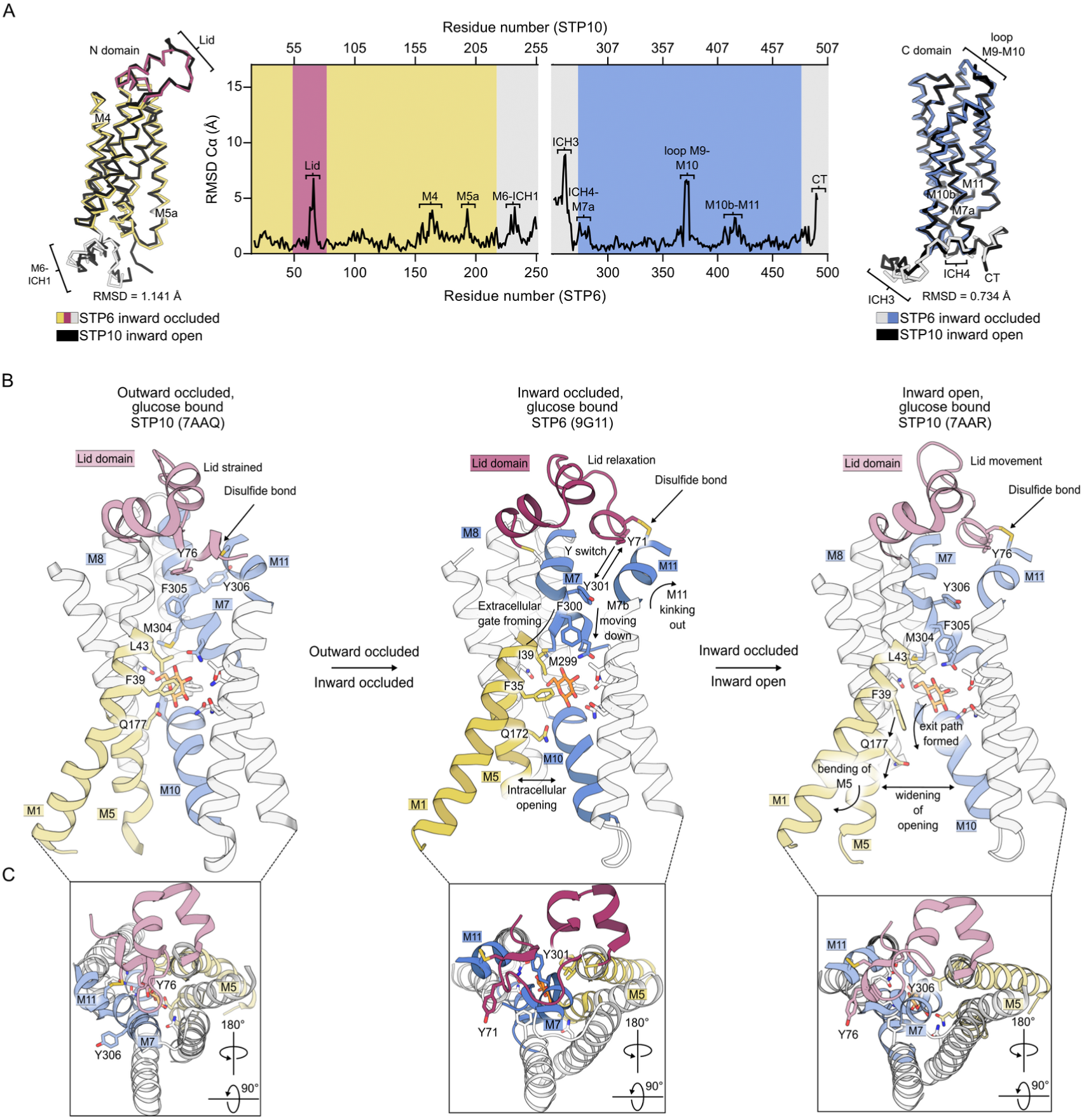
Transporter dynamics transitioning from inward occluded to inward open. **A)** C*α* backbone superposition and the RMSD plotted as a function of backbone position between the inward occluded (STP6, colored) and inward open (STP10, black) states of the N domain (STP6: 17-250 to STP10: 22-255) and C domain (STP6: 251-491 to STP10: 258-498). Regions of interest are highlighted on the ribbon representations of each domain. **B)** Structural changes in the Lid domain (shown in close-up in panel C) and central helical bundle happening upon progression from the outward occluded to the inward open state. Regions of interest are highlighted with colors, pale for STP10 and bright for STP6. Arrows are used to indicate movement and relevant residues are shown as sticks. **C)** Close-up view of rearrangements happening in the Lid domain during progression in the transport cycle from outward occluded to inward open. Regions and residues of interest are annotated.

Simultaneously, a mimetic residue swap between Tyr71 (L3) and Tyr301 (M7b) takes place as part of the dynamic hydrophobic gate controlling access to the binding site (Figure 2B and Supplementary Figure 4), as previously described for STP10 (18). As indicated by the RMSD plot the transporter is seen to undergo subtle rearrangements when transitioning from the inward occluded to the inward open state, with structural deviations being most pronounced in the flexible regions of the C domain, including the loop between M9 and M10 and the C terminal (Figure 2A). Changes are also noticeable in helical regions, including movement of ICH3 and the joint movement of M10b and M11 (Figure 2A and B). When looking at changes happening in residue 0 – 250 the most pronounced deviation results from rearrangement of the Lid domain, which curls away from the central cavity during the transition from the inward occluded to the inward open state (Figure 2A, B and C). The positioning of Tyr71 and Tyr306 is unaffected in this process (Figure 2B and C), suggesting a continuous restriction of the binding site towards the extracellular side. Further movement is found in the ICH1-connected TM region spanning M4-M6, which includes the noticeable outwards kinking of M5a and rearrangement of the polar interactions between the SP and A motifs (Figure 2A, B and Supplementary Figure 3). Accompanied by the large conformational changes in the Lid domain, movement of the TM helices lining the central cavity determines access to the binding site; M1, M2 and M7 come together to secure the binding site towards the extracellular side, going from outward occluded to inward occluded, and kept in the inward open state (Figure 2C and Supplementary Figure 4). Towards the intracellular side the inward transition extends the distance between M4, M5, M10 and M11, successfully creating a path for substrate release (Figure 2B and Supplementary Figure 4). This expansion is initially minor, as the transporter transitions from the outward occluded to the inward occluded state, but dramatically increases to ∼8 Å as it progresses to the inward open state (Figure 2B and Supplementary Figure 4). Thus, the inward occluded conformation of STP6 extends the structural basis for analyzing transporter dynamics of the STPs, while fitting into the current transition scheme described for STP10.

When comparing the *in vitro* transport activities of STP6 and STP10 in identical solid supported membrane (SSM) electrophysiology setups, STP6 and STP10 display comparable affinities towards glucose, with STP6 having a EC_50_ of 452 µM and STP10 a K_M_^app^ of 637 µM (Figure 3A and Supplementary Figure 5). The optimum pH for transport in SSM electrophysiology is pH 5-6, which corresponds to values observed in yeast uptake assays (Supplementary Figures 2 and 5) (17, 19).

**Figure 3.**
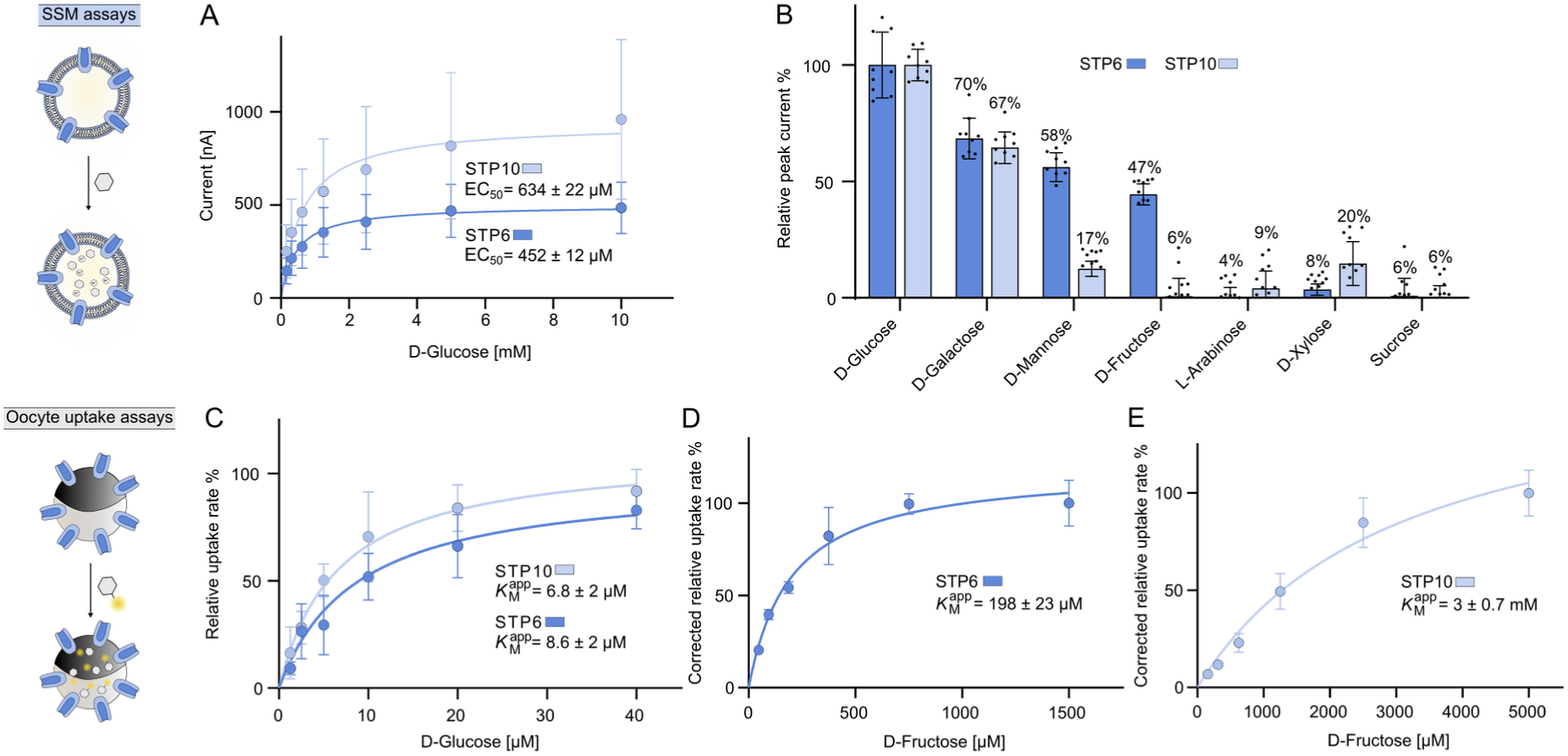
*In vivo* and *in vitro* comparison of STP6 and STP10 transport characteristics. A) Michaelis-Menten analysis of peak currents reported as EC_50_, measured by SSM-based electrophysiology on STP6 and STP10 reconstituted in proteoliposomes when titrated with D-glucose. **B)** Substrate specificity comparison of STP6 and STP10 measured by peak currents conducted from SSM-based electrophysiology when subjected to 10 mM of a range of sugars. Datapoints represent peak currents measured for each substrate at a concentration of 10 mM. Datapoints represent the mean ± SD of n = 3 experiments. *P-* values shown were obtained by one-way ANOVA analysis. SSM experiments were conducted at a symmetrical pH of 7.4 **C)** Michaelis-Menten analysis of labelled D-glucose uptake by STP6 and STP10 determined by heterologous expression in *Xenopus* oocytes. **D)** Michaelis-Menten analysis of labelled D-fructose uptake by STP6 in *Xenopus* oocytes. **E)** Michaelis-Menten analysis of labelled D-fructose uptake by STP10 in *Xenopus* oocytes.

Notably, the EC_50_ measured in this setup represent ∼20-30-fold weaker affinities than the apparent K_M_ value determined in the yeast uptake assays described above. We next tested the SSM-signal towards the substrates described above, including L-arabinose. Here, both STP6 and STP10 showed activity in the presence of glucose, galactose, and mannose but negligible signal in the presence of sucrose and arabinose. In addition, STP6 showed a significant response to fructose, while almost no response was observed for STP10 (Figure 3B). Except for the transport signal observed for xylose with STP10 (Figure 3B), these results match the competition profiles from the yeast uptake assays (17, 19).

To address the affinity differences between the yeast uptake assay and the proteoliposome-based SSM assays, we performed an additional *in vivo* comparison using *Xenopus laevis* oocytes. In oocytes, STP6 and STP10 display affinities for glucose transport that are 2-3-fold stronger than those observed in yeast uptake assays, with respected K_M_^app^ of 8.6 µM and 6.8 µM (Figure 3C and Supplementary Figure 6). Moreover, alanine mutation of the suggested proton binding site in STP6 (Asp38) and the corresponding site in STP10 (Asp42) abolish uptake of glucose in oocytes, underlining the fundamental role of this residue in coupling sugar transport to the PMF (Supplementary Figure 6).

Next, we used radiolabeled fructose to quantify transport in STP6 and STP10. Here, fructose transport was observed for both transporters, however, while STP6 display intermediate affinity, (198 µM, ∼20-fold weaker than for glucose) a considerably lower fructose affinity is observed for STP10 (K_M_^app^ = 3000 µM, ∼440-fold weaker than for glucose) (Figure 3D, E and Supplementary Figure 6). The drastically lower fructose affinity of STP10 compared to STP6 correlates well with the differences observed in competition assays and response currents measured in proteoliposomes.

While fructose transport has previously been suggested for STP6 (9), the observed differences in response to fructose between STP6 and STP10 is surprising given the virtually identical binding sites (Supplementary Figure 8). Therefore, we looked at how the global structural rearrangements happening during transport affect the local environment of the substrate, by comparing the binding pocket in structures of STP6 and STP10. When superposing the C domains of STP6 and the outward occluded state of STP10, a clear displacement of Phe35 and Gln172 is apparent, which is directly linked to the rearrangement of M1 and M5 (Figure 4A Panel 1 and Figure 2). Interaction with Phe35 and Gln172 is known to be crucial for high-affinity glucose transport (18) and the displacement thus results in a looser binding pose, effectively lowering the affinity when transitioning to the inward state. The glucose molecule remains bound as escape is not possible in the inward occluded state.

**Figure 4.**
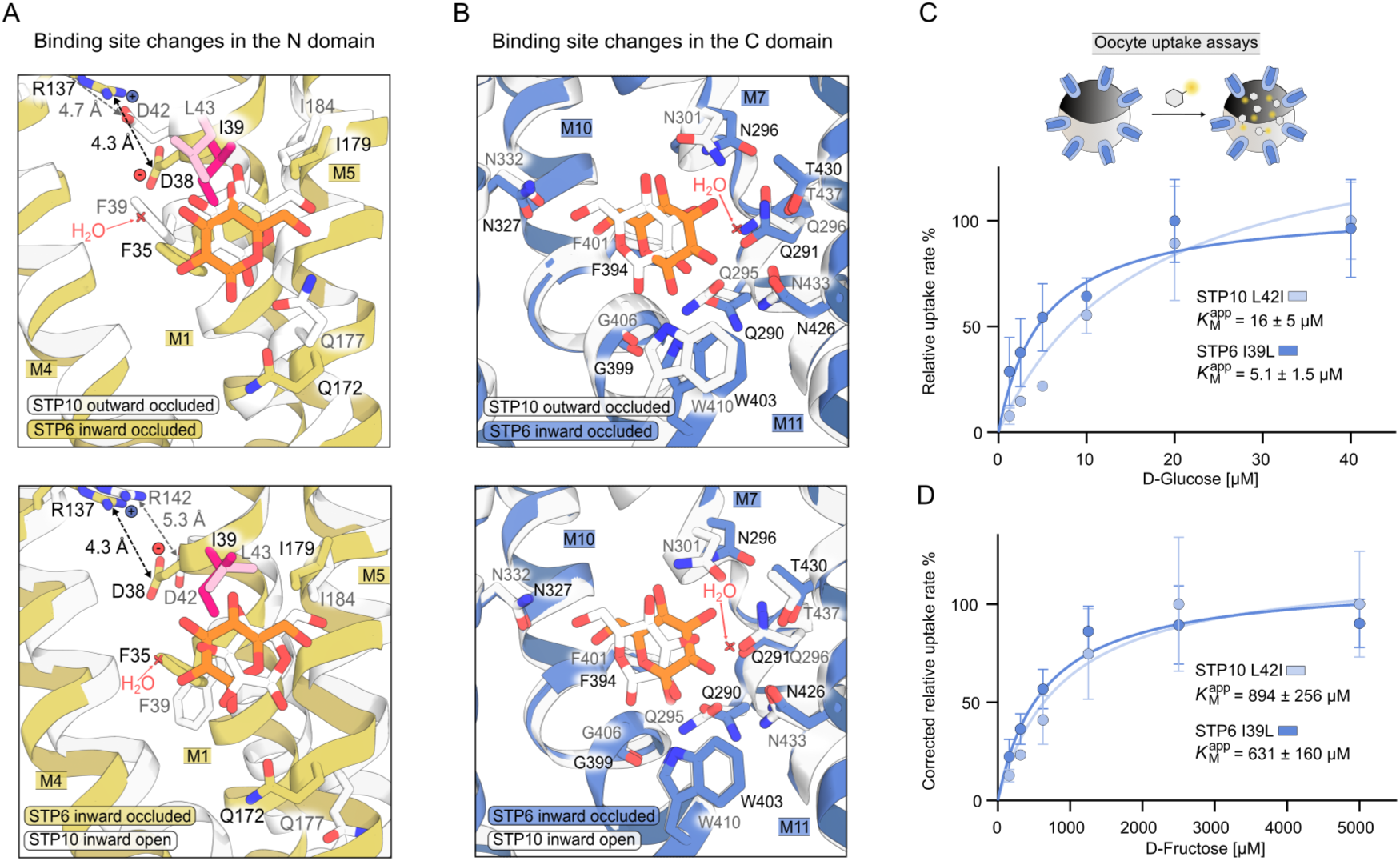
Structural and functional analysis of the binding sites in STP6 and STP10. **A)** Comparison of binding pocket interactions found in the N between the outward occluded and inward open STP10 states and the inward occluded STP6 state. The only differing residues I39 (STP6) and L42 (STP10), are highlighted in hot and light pink, respectively. **B)** Comparison of binding pocket interactions found in the C domain. For both C and D structural superposition has been performed on the C domain. STP10 structures are shown in white, while STP6 is colored (N domain in yellow and C domain in blue). The depicted water molecules belong to the STP10 structures. **C)** Michaelis-Menten analysis on labelled D-glucose uptake by binding site switch mutants, STP6 I39L and STP10 L43I in *Xenopus* oocytes. **D)** Michaelis-Menten analysis on labelled D-fructose uptake by binding site switch mutants, STP6 I39L and STP10 L43I in *Xenopus* oocytes. Datapoints represent the mean ± SD of n = 4-6 biologically independent experiments. *Xenopus* oocyte experiments were performed at pH 5.

Progression from the inward occluded to the inward open state confers further displacement of Phe35 (M1) and Gln172 (M5), observed to be fully disengaged from the glucose, completing an exit path and allowing for substrate release (Figure 4A Panel 2, Figure 2B and Supplementary Figure 4). In contrast to the observed changes in the N domain, substrate interactions with the C domain remain unchanged throughout the transition (Figure 4B). Thus, the majority of the local structural rearrangements happening upon transporter transitioning take place in the N domain. The 5 Å distance between Asp38 and Arg137 observed in STP6 (Figure 4A) supports protonation of Asp38, meaning that the STP6 structure represents a proton-loaded state (18). The only noticeable difference is observed at the top of the binding site where Ile39 in STP6 corresponds to Leu43 in STP10, resulting in the positional change of a single methyl group (Figure 4A, B and Supplementary Figure 7).

To explore if this subtle change had an effect on the transport properties of STP6 and STP10, we used point mutations to recreate the binding site of STP10 in STP6, essentially shifting the methyl group by one carbon atom. This mutant, STP6 I39L, displays a 3-fold reduction in fructose affinity (K_M_^app^_STP6 I39L_ = 631 µM vs. K_M_^app^_STP6 (WT)_ =198 µM), while slightly improving affinity towards glucose (K_M_^app^_STP6 I39L_ = 5.1 µM vs. K_M_^app^_STP6 (WT)_ = 8.6 µM) (Figure 4C and D), reflecting the glucose affinity of WT STP10. The inverse trend was found in the reverse experiment, recreating the binding site of STP6 in STP10, by introducing the mutation L43I. STP10 L43I shows a 3-fold improved affinity for fructose (K_M_^app^_STP10 L43I_ = 894 µM vs. K_M_^app^_STP10 (WT)_ = 3000 µM) while glucose affinity is slightly reduced (∼2-3-fold K_M_^app^_STP10 L43I_ = 16 µM vs. K_M_^app^_STP10 (WT)_ = 6.8 µM) (Figure 4C and D). This result shows that the ∼2 Å shift of a single methyl group in the binding site is sufficient to modulate glucose and fructose specificity in Sugar Transport Proteins.

## DISCUSSION

Here we characterize and compare *A. thaliana* STP6 and STP10, and present a 3.2 Å crystal structure of STP6 with glucose bound. The STP6 structure captures a previously unresolved "inward occluded state" of the STP transport cycle. This state has previously been resolved only for the bacterial homolog XylE (29, 41). By expanding our structural knowledge with the inward occluded state of STP6, we have gained new insight into the dynamics governing substrate release in the STP transport cycle. The STP structures collectively point to a model where substrate release takes place in two consecutive steps: First, global rearrangement of the Lid domain and helical regions brings the transporter into the inward facing orientation. Next, the central cavity of the transporter is expanded towards the extracellular side, shaping the substrate exit path. In this process a hydrophobic extracellular gate is formed by Ile39, Met299 and Phe300, while Phe35 (M1) and Gln172 (M5) are retracted from the binding pocket, enforcing substrate release by lowering the local affinity (Figure 2B). The interaction observed between CHS and Arg340 (M8) in the C domain A-motif of STP6 likely stabilizes the transporter in the inward conformation, suggesting that the immediate environment may be involved in regulation of the conformational equilibrium. This idea aligns with findings from other studies on SP family members and ties into a more general discussion of how transporters are affected by membrane composition (31, 39, 42–44).

Considering the effects of external factors is important when assessing transport properties in biophysical assays. Challenges with quantifying protein levels lead to a lack of meaningful Vmax values for membrane protein transporters, limiting affinity assessments to apparent affinities, i.e. K_M_-values (45).

In the analysis of transport kinetics we have used three different experimental methodologies (yeast uptake, oocyte uptake and SSM) to assess kinetic parameters for STP6 and STP10. While we use apparent K_M_-values for *in vivo* analysis, we denote the apparent affinity derived from SSM as EC_50_, because the peak currents can contain a mixture of binding and transport signal that cannot always be deconvoluted (46, 47). We demonstrate a significant and systematic deviation between methodologies, most pronounced between SSM and the *in vivo* based experimental setups (∼20-100 fold), due to these overlapping electrogenic events in SSM. But there is also noticeable difference between *in vivo* setups (2-10 fold). Factors expected to influence kinetics of transport include buffer composition, lipid composition of the membrane, and the proton motive force, consequently giving rise to observable systematic differences between methodologies (48–50). These factors are especially present in SSM, as experiments are conducted in absence of a pH gradient, and transport is driven only by the electrochemical gradient of the substrate (51). For these reasons, SSM analysis often gives rise to weaker apparent affinities compared to *in vivo* based systems. Our work demonstrates the necessity for caution when comparing transport properties between proteins across methodologies, and identical experimental setups should ideally be used when possible.

In our side-by-side comparison of STP6 and STP10, we investigate how STPs discriminate between substrates. We find a 15-fold difference in fructose affinity between WT STP10 and WT STP6 in oocyte uptake assays. Next, we show that we can modulate fructose accommodation by changing the position of a single methyl group forming part of the extracellular gate, resulting in a gain of affinity for fructose, as seen for STP10 L43I. This notable gain of fructose affinity results in only minor decrease of glucose affinity, supporting the general ligand interaction concept of high affinity imposing high specificity (52). Changing the equivalent position in STP6 similarly led to a notable decreased fructose affinity and only minor increased glucose affinity.

These results correspond well with our previous specificity-affinity concept suggested for the STPs, where the polar network of the C domain confers hexose specificity, showing here that affinity differences between different hexoses arises from hydrophobic interactions with the N domain (17, 18). Furthermore, STP8 and STP13 previously shown to transport fructose also contain an isoleucine on the position equivalent to I39 in STP6 and another study on a homolog from green algae revealed that the binding site position equivalent to Ile39/Leu43, could be used to modulate affinities of glucose and galactose in a similar manner to here (34, 37, 53). This suggests a conserved mechanism for controlling affinity induced specificity by transporters belonging to the SP family, which we can now expand on. Substrate interactions mediated by residues localized in the N domain control affinity by direct interactions, as seen for Phe35 (M1) and Gln172 (M5) (18), or indirectly by shaping and tightening of the binding pocket, as seen for Ile39 (M1) and Met299 (M7). As demonstrated here, indirect interactions are highly sensitive and minute changes can be used to modulate the affinity-induced specificity. For fructose binding, we hypothesize a binding scheme closely mimicking the one displayed by glucose. Fructose is found in a pyranose form and furanose form in a ∼2:1 equilibrium in solution, as opposed to glucose which is principally only found on the pyranose form (99%) (54–56). Therefore, we speculate that for fructose, the pyranose form is selectively transported over the furanose form by STPs.

In summary, we have shown that changing the position of a single methyl group is sufficient to manipulate substrate affinities of STPs, showing that subtle variation in the hydrophobic binding site is a key regulator of selectivity. The less-than-perfect fructose affinity complementation of STP6 I39L and STP10 L43I to their respective WT counterparts implies the presence of other yet undiscovered regulatory elements and/or kinetic influence by other regions of the protein (57, 58). In STPs, the extracellular gate formed between M1 and M7 involves Ile39/Leu43 and constitute hydrophobic interactions (Figure 2B). Notably, this gate is formed by polar interactions in other SP proteins, and has been found to be central to the conformational change from the outward to the inward state (58). This suggests that the extracellular gate might play a broader role in shaping substrate specificity.

Another interesting feature of the STPs is the Lid domain. Given its exofacial localization and great sequence variability (Supplementary Figure 8), it is tempting to speculate on a potential role as a regulatory element, which may give rise to unique transport properties or substrate promiscuity. For instance, recent studies have shown disaccharide transport by STPs found in crops (10, 16, 59). This shows that while we have established a strong understanding of how monosaccharides interactions are mediated by STPs, there is still much to uncover about potential regulatory elements and substrate promiscuity. Uncovering how this is achieved and how it relates to specific stages of plant development will be valuable and can provide a foundation for plant engineering in the future.

## Acknowledgements

The authors acknowledge beamlines I24 and I04 at the Diamond Light Source and beamline BioMAX at the MAX IV Laboratory, where X-ray data were collected, as well as DESY-PETRA III for crystal screening. We acknowledge access to the computational infrastructure at the Center for Structural Biology at Aarhus University. This project has received funding from the European Research Council (ERC) under the European Union’s Horizon 2020 research and innovation programme (grant agreement No 101000936) and Novo Nordisk Foundation grant NNF24OC0088380 to BPP. AB was supported by Fellowship from Scholars at Risk from Ukrainian Universities.

## Author Contributions

Sample preparation: CA, LB

Structural data collection and analysis: CA, LB, BPP

Activity assays: CA, LB, AB, AC, SB, JHD

Manuscript preparation: CA, BPP

Figure preparation: CA, LB

## Data and materials Availability

Atomic model and structure factors for STP6 have been deposited in the Protein Data Bank (PDB) with the accession code 9G11.

## Materials and Methods

### Protein expression and purification

The gene encoding *A. thaliana* STP6 (UniProt accession number Q9SFG0) was cloned into an p423_GAL1 expression vector containing a C-terminal purification tag with a thrombin cleavage site followed by a 10x histidine tag. Protein expression was performed using *S. cerevisiae* (strain DSY-5) grown in a culture vessel. High cell density was obtained by exponential fed-batch, induced with galactose, and harvested after ∼18 hours (70). Harvested cells were washed in cold water, spun down and resuspended in lysis buffer (600 mM NaCl, 100 mM Tris-HCl pH 7.5 and 1.2 mM PMSF), followed by opening of cells using beating in cooled bead canisters filled with 0.5 mm glass beads.

Opened cells were centrifuged for 20 min, 17.000 RCF at 4 °C, followed by isolation of membranes by ultracentrifugation of the supernatant at 200.000 RCF for 2 h. Pelleted membranes were homogenized in membrane buffer (500 mM NaCl, 50 mM Tris-HCl pH 7.5, 20% (w/v) glycerol) and flash frozen in liquid nitrogen. Protein purification was carried out with 6 g of thawed membranes, solubilized for 30 minutes in solubilization buffer (150 mM NaCl, 50 mM Tris pH 7.5, 5% glycerol, 1% n-dodecyl-β-D-maltoside (DDM) and 0.1% cholesterol hemisuccinate (CHS)) in a total volume of 100 mL. Unsolubilized material was removed by centrifugation followed by filtration of the supernatant using a 1.2 µM filter. 20 mM imidazole pH 7.5 was added prior to loading solubilized membranes on a pre-equilibrated 1 mL Ni-NTA column (GE Healthcare) at 1 mL min^-1^. The column was washed with 50 column volumes of W60 buffer (150 mM NaCl, 50 mM Tris pH 7.5, 0.1% DDM, 0.01% CHS, and 60 mM imidazole pH 7.5), followed by washing 30 column volumes in G20-buffer (20 mM Tris-HCl, 200 mM NaCl, 20 mM imidazole pH 7.5, 0.12% octyl glucose neopentyl glycol (OGNG), 0.012% CHS or 0.02% DDM, 0.002% CHS). Bound protein was cleaved from the column by circulating in 5 mL G-buffer supplemented with 200 U of bovine thrombin for ∼14 hours at 19 oC. A final wash was performed using 15 mL of G-buffer supplemented with 40 mM imidazole pH 7.5.

The sample was concentrated using 50 kDa MWCO spin column (Merck) to a final volume of ∼450 µL prior to size-exclusion chromatography performed on Bio-Rad system equipped with a Superdex200inc. column (GE Healthcare), equilibrated in G-buffer optimized by a thermostability assay (71) (20 mM Tris-HCl, 200 mM NaCl, 0.12% OGNG, 0.012% CHS or 0.02% DDM, 0.002% CHS). The choice of detergent dependent on downstream protein application. OGNG was used in crystallization assays whereas DDM was used for proteoliposome reconstitution.

### Yeast uptake assay

Functional characterization in yeast was performed by introducing the A. thaliana STP6 gene into the p426_MET25 vector, allowing constitutive expression under a methionine-inhibited promoter. Transformation was carried out into S. cerevisiae lacking endogenous hexose transporters (strain EBY.WV4000) (60) and plated on SDA plates without uracil supplemented with 2% maltose. After three days a streak of cells was inoculated into 50 mL of complementary media and grown to an OD600 of ∼1.5. Cells were washed with 25 mM NaPO4 buffer pH 5.0, followed by centrifugation and resuspension, achieving a final OD600 of 10. Experiments were carried out using 20 µL of fresh cells diluted to a final volume of 200 µL with NaPO4 buffer (pH directed by the experiment). Reactions were started upon addition of substrate containing 1 µCi of radioactively [3H]-labelling. Time-course experiments with glucose were performed with 2 – 640 µM glucose on STP6 and empty plasmid which was subsequently subtracted as background (noted as corrected). Time-course experiments were then treated as end-point measurements for each time point and the resulting apparent KM values were determined (Supplementary figure 2), to justify the use of end-point measurements in other experiments. For additional experiments the following concentrations of substrate was used: 100 µM was used in time dependent assays with galactose and mannose and 10 µM glucose was used in pH dependent and competition assays. Competing sugars were added in 25x excess and for inhibitors 500 µM was used, except for CCCP where 100 µM was used. All inhibitors were dissolved in DMSO. Reactions were stopped in noted intervals by collecting cells on a cellulose filter (0.8 µm) under vacuum. Three wash cycles were then performed before retracting each filter into individual scintillation tubes and addition of 3 mL of scintillation liquid. Quantification of [3H]-signal was done using Tri-Carb 5110TR Liquid Scintillation Counter (Perkin Elmer). Data analysis was performed in GraphPad Prism 9 where linear regression (time dependent uptake) or Michaelis-Menten fit (concentration dependent uptake) was utilized and afterwards normalized.

### Solid-supported membrane (SSM)-based electrophysiology assays

SSM-based electrophysiology experiments were performed by reconstituting purified STP6 and STP10 into proteoliposomes analyzed utilizing a SURFE2R N1 instrument (Nanion Technologies). For all experiments sensors preparation was performed as described in (47). Proteoliposomes were prepared using desiccated yeast polar lipid extract (Avanti), resuspended in surfer buffer (30 mM HEPES pH 7.4, 140 mM NaCl and 5 mM MgCl2). Liposomes were homogenized in size by extrusion through a 0.4 µM membrane followed by pulse sonication in water batch for 1 min. Liposomes were destabilized by addition of 1% (v/v) n-octyl-β-D-glucoside prior to mixing with purified protein, achieving a final LPR=5, or with addition of buffer in equal volume (control). Detergent was extracted by stirring suspensions overnight with 400 mg mL-1 BioBeads (Bio-Rad) at 4 °C. Proteoliposomes were diluted 1:5 in surfer buffer prior to application onto 3 mM sensors. Sensors were centrifuged at 2100 g for 30 min. Activating and non-activating buffers were prepared as the surfer buffer, however, the activating buffer was supplemented with 0-40 mM of D-glucose (concentration dependent assays) or 10 mM of the analyzed sugar (substrate specificity assays). A single solution exchange workflow was used for all experiments. Michaelis-Menten curves were fitted using the peak currents (containing binding and transport signal), datapoints represent the mean of three individual experiments and the SD represents deviation across all individual data points. Data handling was performed in GraphPad prism. As binding and transport signal inseparable we denote the derived kinetic constants as EC_50_ instead of apparent *K*_M_.

### *Xenopus* Oocyte assays

Uptake assays in *Xenopus* oocytes were performed as described in (61) and (62), with few modifications. The gene encoding A. thaliana STP6, STP10 and mutagenic versions, created with Q5 site-directed mutagenesis (NEB), were encoded into the pNB1-U vector. Template for in vitro cRNA transcription was amplified using primers annealing to the 5’ and 3’ UTR implementing a poly(A)-tail (FW-HNN49: AATTAACCCTCACTAAAGGGTTGTAATACGACTCACTATAGGG and RV-HNN50: TTTTTTTTTTTTTTTTTTTTTTTTTTTTTATACTCAAGCTAGCCTCGAG). In vitro transcription was carried out with the mMESSAGE mMACHINE T7 Transcription Kit (Thermo Fisher) using 200 ng of template. Adult *X. laevis* oocytes from multiple females (EcoCyte Bioscience) were injected with 25 ng of cRNA or RNAse free water (control) using the Nanoject III (Drummond Scientific) equipped with capillary needles. Oocytes were incubated at 16 °C for 3 days in MBS buffer pH 7 supplemented with 10 IU/mL gentamycin. Time-course experiments using 1.25-40 µM D-glucose supplemented with 1 µCi/mL [3H]-D-Glucose, where performed to derive Michaelis-Menten kinetics for WT STP6. Additionally, time-course experiments were treated as end-point measurements and the resulting apparent KM-values were determined (Supplementary figure 6), justifying the use of end-point measurements in other experiments. Additional uptake assays were performed as end-point measurements with 1.25-40 µM D-glucose supplemented with 1 µCi/mL [3H]-D-Glucose or 0.16-5 mM of D-fructose supplemented with 5 µCi/mL [14C]-D-fructose in ND-96 buffer (96 mM NaCl, 1 mM MgCl2, 2 mM KCl, 1.8 mM CaCl2, 5 mM MES pH 5). Oocytes were pH-equilibrated for 5 min in ND-96 pH 5 buffer before addition of substrate. Experiments were run for 30 min unless otherwise stated. To stop the reactions oocytes were individually extracted and washed three times in excess ND-96 buffer, before transferring into scintillation vials. The oocytes were lysed by addition of 200 µL 10% SDS, followed by sonication in sonicator bath. 3 mL of scintillation liquid was added before quantification of radioactivity using Tri-Carb 5110TR Liquid Scintillation Counter (Perkin Elmer). Each oocyte was handled as a unique experiment and each datapoint represents 3-5 oocytes. Data analysis was performed in GraphPad Prism 9. For each hot solution, the efficiency was measured after each experiment using 100 µL of buffer. The efficiency and the specific activity of the diluted substrate was used to calculate uptake for each oocyte in pmol, following the example in (63). Disintegrations per minute (DPM) was adjusted according to the measured efficiency, allowing to determine the total amount of Ci taken up over time (DPM/2,22*10^12). The specific activity was used to determine the uptake in pmol for each oocyte. The data was then normalized to the relative Vmax before fitting of Michaelis-Menten kinetics (concentration dependent uptake) or linear regression (time dependent uptake). Assays performed with fructose were corrected for background signal using water injected oocytes, noted as “corrected” on the legend of the Y-axis.

### Crystallization

Peak fractions from size-exclusion chromatography were concentrated to 4-6 mg mL-1 using a 50 kDa MWCO spin column (Merck), followed by addition of 30 mM D-glucose. Crystallization was carried out as vapor diffusion with sitting drops in MRC 2-well plates and MRC MAXI (SWISSCI) set up with the Mosquito lipid handler (TTP Labtech). A protein:reservoir ratio of 1:1 was used with the initial drop size of 400 nL later scaled up to 1.2 µL carried out at 19 oC. Crystals appeared after 2-3 days in 28-32% PEG smear (PEG2000, PEG3000, PEG3350, PEG4000 and PEG5000), 0.1 M Hepes, pH 8.0 and 0.2-0.5 M NaCl. The final dataset was collected at Diamond Light Source I24 with a wavelength of 0.9686 Å.

### Structural data processing

Datasets were processed in XDS followed by quality assessment in phenix.xtriage suggesting crystal twinning. Space group validation was carried out in Zanuda (64) and P3221 was given as the most likely space group, with merohedral twin operators (-h,-h,l) and one copy in the asymmetric unit (∼77% solvent content). Final data-scaling was carried out in AIMLESS (65) giving a resolution range of 84-3.2 Å. The phase problem was solved using Molecular Replacement in phenix.phaser (66) with the inward open structure of STP10 (PDB 7AAR), prepared with Sculptor, as search model.

Manual model building was done in Coot (67) with iterative model adjustment in NAMDINATOR (72) and model refinement in phenix.refine (68) with final refinement parameters including individual ADP weighting and grouped TLS (2 groups). The final model has an R-work of 22.65% and an R-free of 25.16%. Structure geometry was validated by monitoring MolProbity (69) and Ramachandran score (73). Statistics of the model refinement can be found in Supplementary Table 1. Figures were prepared using PyMOL Molecular Graphics System (Schrödinger, LLC) (74). Conservation of residues across species was analyzed using Consurf (75). Sequence alignments were constructed with PROMALS3D (76)

## SUPPLEMENTARY FIGURES LEGENDS

**Supplementary Figure 1.**
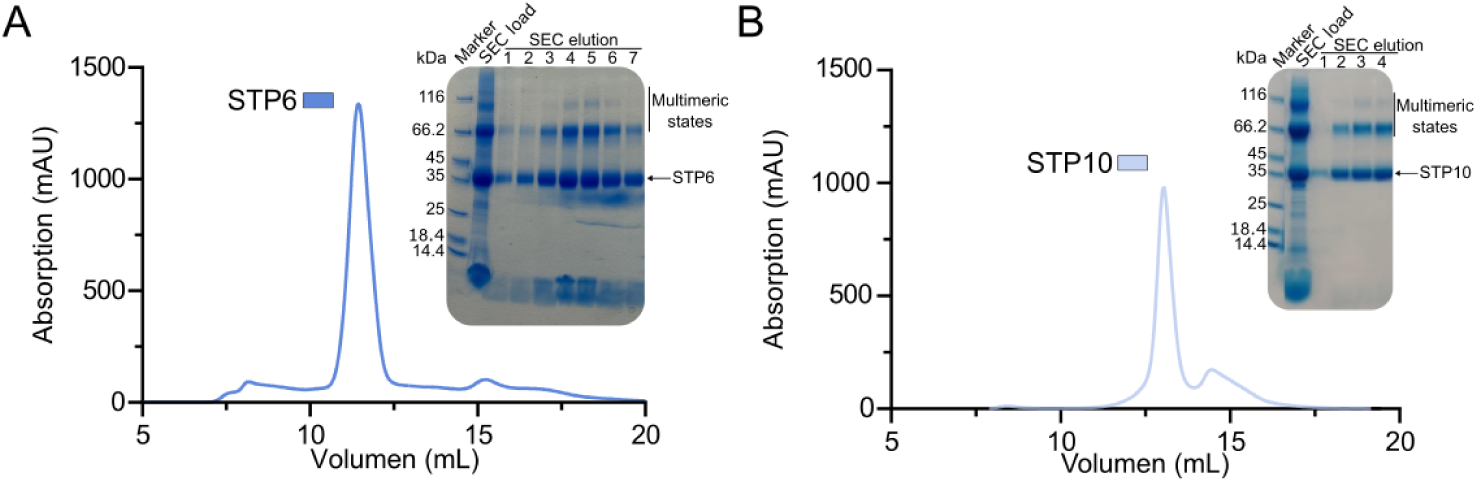
Initial expression and purification of STP6 and STP10. A) Size-exclusion chromatography profile of STP6 with representative SDS-PAGE depicting protein purity. B) Size-exclusion chromatography profile of STP10 with representative SDS-PAGE depicting protein purity. Expression and purification of both targets have been carried out multiple times with reproducible results.

**Supplementary Figure 2.**
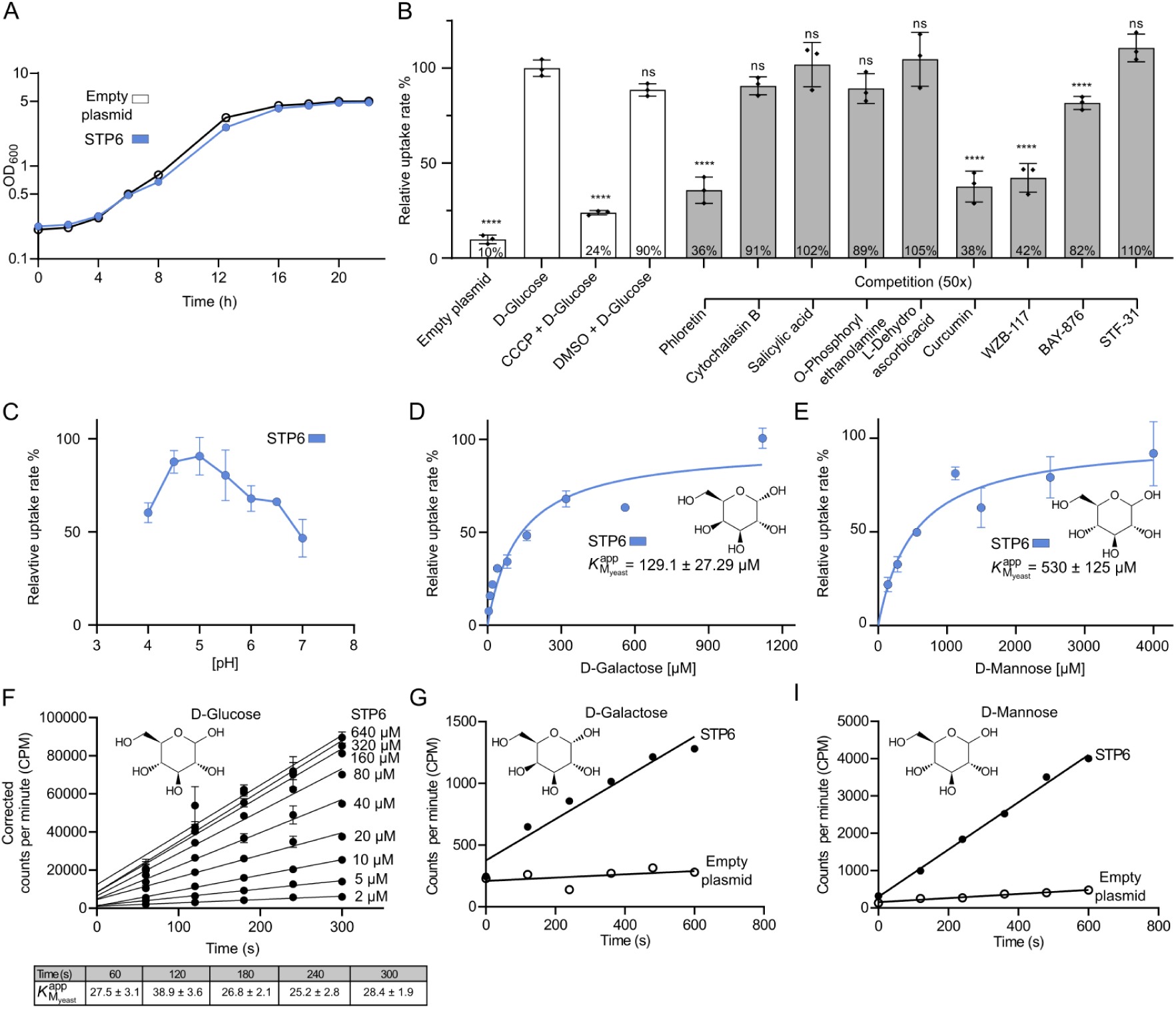
Functional characterization of STP6 in yeast uptake assays. **A)** Growth of EBY.VW4000 expressing either empty plasmid or STP6. Growth was monitored by OD_600_ of three independent cultures over 22 hours. The exponential phase is visible from OD_600_ 0.4-4. The Y-axis is shown on a logarithmic scale (log10). **B)** Glucose competition assay on STP6 with a range of potential inhibitors performed in yeast cells using 10 µM of D-glucose and 500 µM of each of the tested inhibitors, expect for CCCP where 100 µM was used. Datapoints represent mean *±* SD of n = 3 replicate experiments. *P*-values for the competition assay were calculated by one-way ANOVA followed by Dunnett’s multiple comparisons test. ****, *p* < 0.0001. **C)** pH dependency of uptake by STP6 performed using 10 µM D-glucose in yeast cells. **D)** Michaelis-Menten analysis of D-galactose uptake by STP6 in yeast cells. **E**) Michaelis-Menten analysis of D-mannose by STP6 in yeast cells. Datapoints represent the mean ± SD of n = 3 experiments. Assays were performed at pH 5 unless stated otherwise. **F)** Time dependent uptake of D-glucose by STP6 over a concentration range from 2 – 640 µM corrected for background signal measured on cells transformed with empty plasmid, used to calculate the apparent *K*_M_ value shown in figure 1, panel A. The table shows apparent *K*_M_ values determined by treating the data as end-point measurements for each of given time points. The variation in *K*_M_ values reflects the observed SD among the individual datapoints used in the analysis shown. **G)** Time dependent uptake of D-galactose by STP6 (black circles) or empty plasmid (empty circles) into yeast cells. **H)** Time dependent uptake of D-mannose by STP6 (black circles) or empty plasmid (empty circles) into yeast cells. All of the above experiments were carried out at pH 5.

**Supplementary Figure 3.**
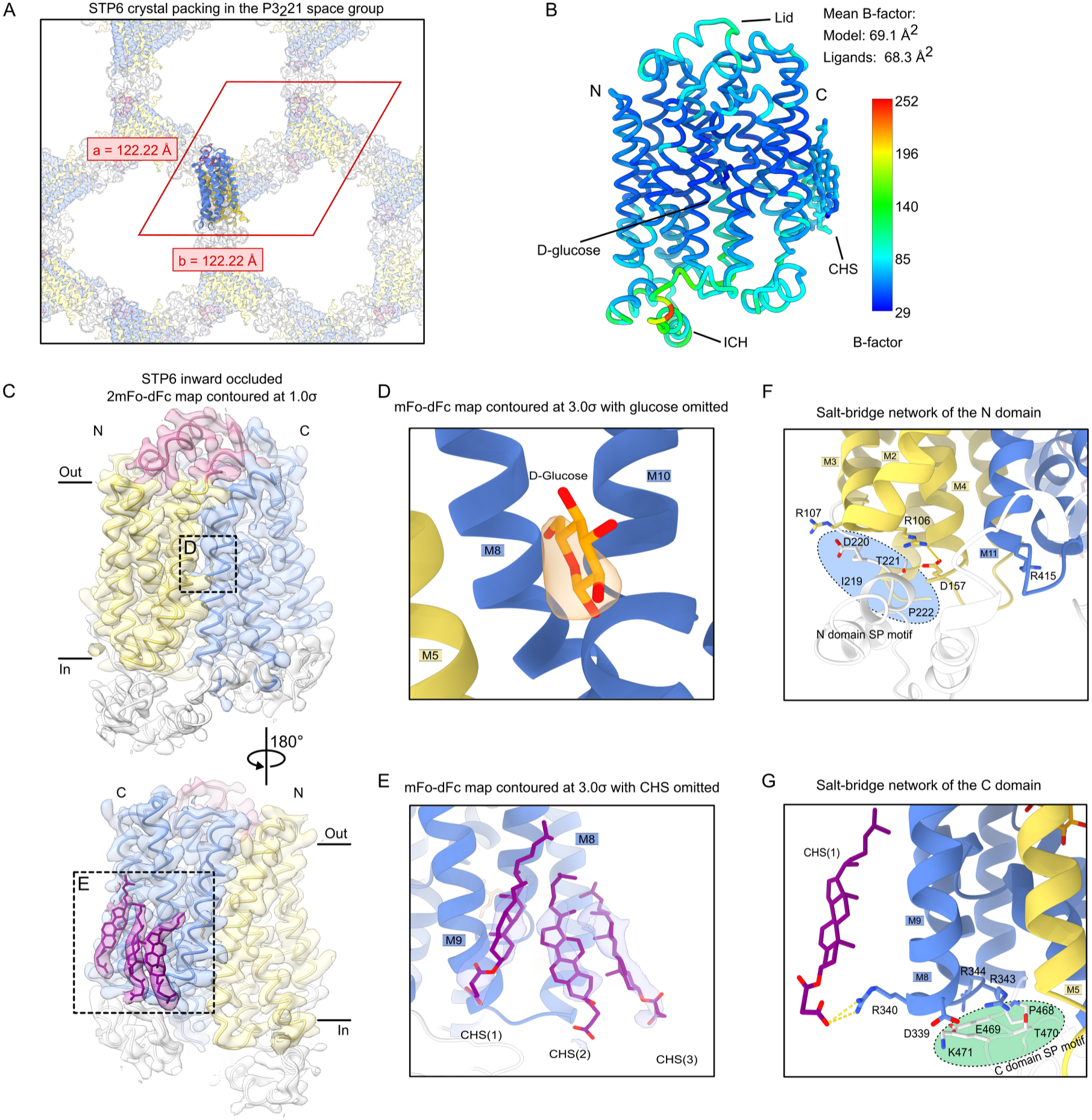
STP6 crystal packing and close-up views of important regions. **A)** The crystal packing of STP6 with the asymmetric unit highlighted by the red rhombus. The a and b axis are shown in red. B) Ribbon representation of the STP6 backbone colored according to the atomic displacement factor (B-factor) with a gradient from blue (29 Å^2^) to red (252 Å^2^). C) The 2mFo-dFc electron density after the final refinement (contoured at 1*σ*) and shown as transparent surface, with the final model overlaid. Ligand bound regions are highlighted in panel D and E. D) The mFo-dFC map (contoured at 3*σ*) calculated after omitting the glucose in the final model. E) The mFo-dFC map (contoured at 3*σ*) calculated after omitting the CHS molecule in the final model. F) Interactions present between the SP and A motifs in the N domain. G) The SP and A motif of the C domain, including a close-up view of the salt-bridge formed between R340 and the CHS head group. The SP motifs are highlighted by ovals. Yellow dashes indicate interactions between residues.

**Supplementary Figure 4.**
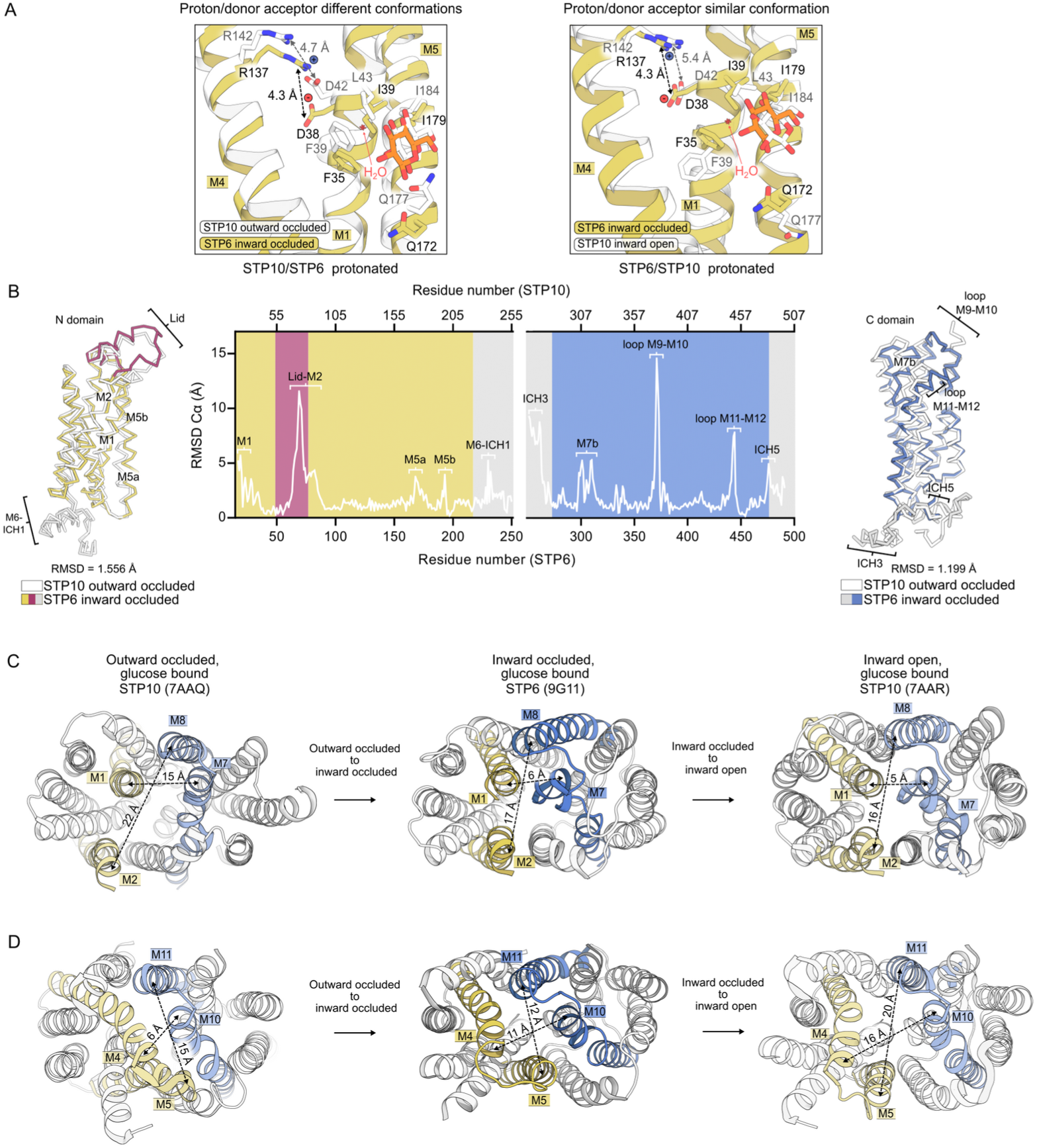
Transporter dynamics during progression through the transport cycle. **A)** Comparison of the proton/donor acceptor pair (D42/142 and D38/R137) sitting in the N domain between the outward occluded and inward open STP10 states and the inward occluded STP6 state. Different orientations are observed between the outward and inward occluded states, while similar orientations are seen in the inward occluded and open state. B) Cα backbone superposition and the RMSD plotted as a function of backbone positioning between the inward occluded (STP6, colored) and outward occluded (STP10, white) states of the N domain (STP6: 17-250 to STP10: 22-255) and C domain (STP6: 251-491 to STP10: 258-498). Regions of interest are highlighted on the ribbon representations of each domain. The STP6 domain coloring follows the schematic shown in the main figures. N domain yellow, Lid domain magenta, ICH domain white and C domain blue. C) Top views showing transporter occlusion towards the extracellular side of the three conformational states, outward occluded, inward occluded and inward open. Cavity lining helices; M1, M2, M7 and M8 are highlighted with colors and the distances between them are shown. D) Bottom views showing transport opening towards the intracellular side during progression from outward occluded to inward open conformational state. Distances between cavity lining helices; M4, M5, M10, M11 (colored) are shown.

**Supplementary Figure 5.**
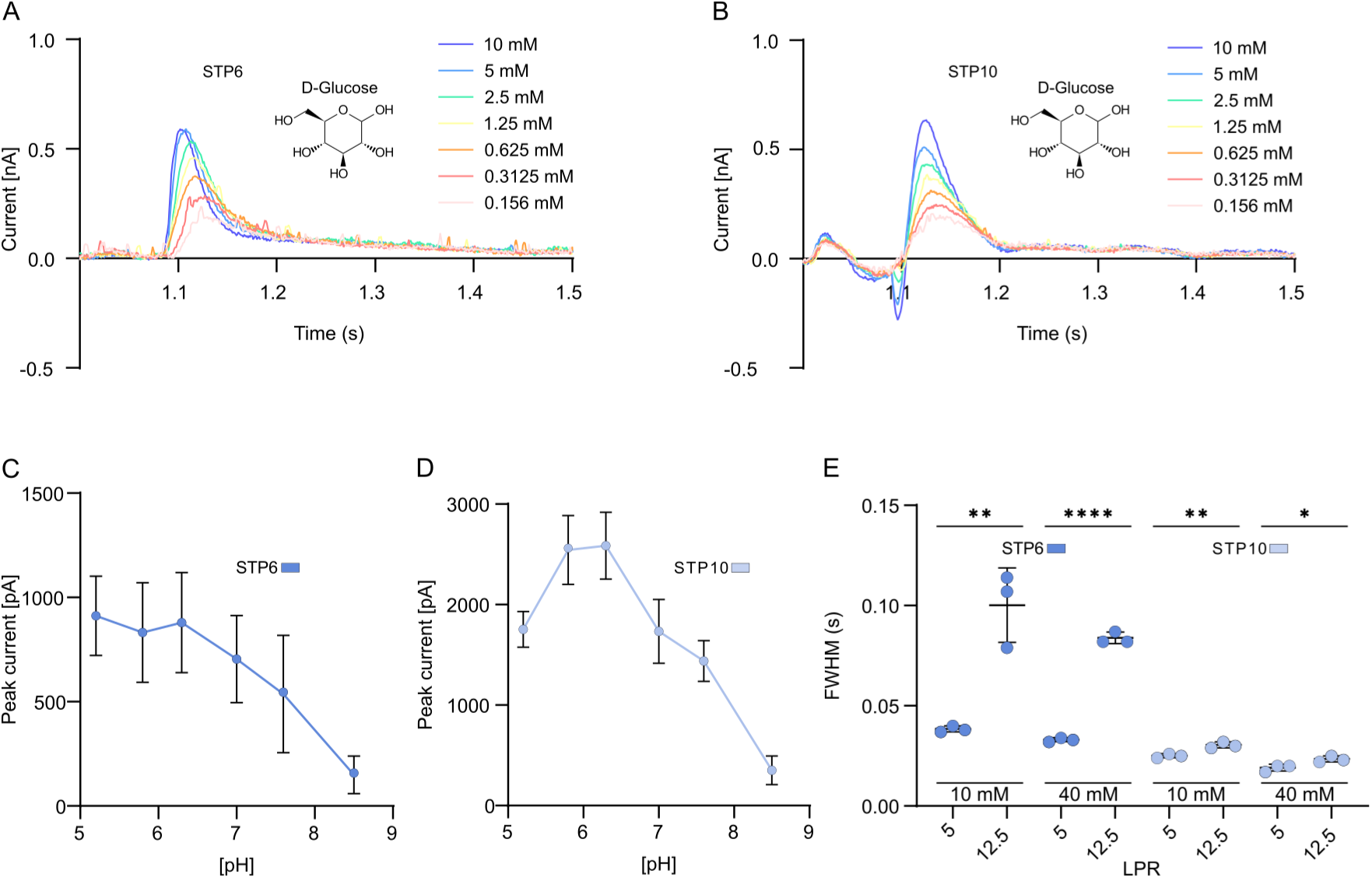
Transport activity of STP6 and STP10 determined by SSM-based electrophysiology. **A)** Raw traces of currents obtained by SSM-based electrophysiology when titrating STP6 containing proteoliposomes with a range of D-glucose (0.156 – 10 mM). **B)** Raw traces of currents obtained when titrating STP10 containing proteoliposomes with a range of D-glucose (0.156 – 10 mM). Traces show data obtained from a representable experiment performed on one sensor with three technical replicates. Experiments were performed at a symmetrical pH of 7.4. **C)** Peak currents elicited by addition of 10 mM glucose to proteoliposomes with STP6 over the indicated symmetrical pH range. Error bars represent ± SD of n = 3 measurements. **D)** Peak currents elicited by addition of 10 mM glucose to proteoliposomes with STP10 over the indicated symmetrical pH range. Error bars represent ± SD of n = 3 measurements. **E)** Change in peak width elicited by 10 or 40 mM glucose at half maximum (FWHM) in relation to changing lipid to protein ratio (LPR), for proteoliposomes containing STP6 and STP10. Error bars represent ± SD of n = 3 measurements. Data was analyzed by unpaired t-tests (*P-*values as follows: ** = *p* = 0.0045-0.0072, **** = *p* < 0.0001, * *p* = 0.0314).

**Supplementary Figure 6.**
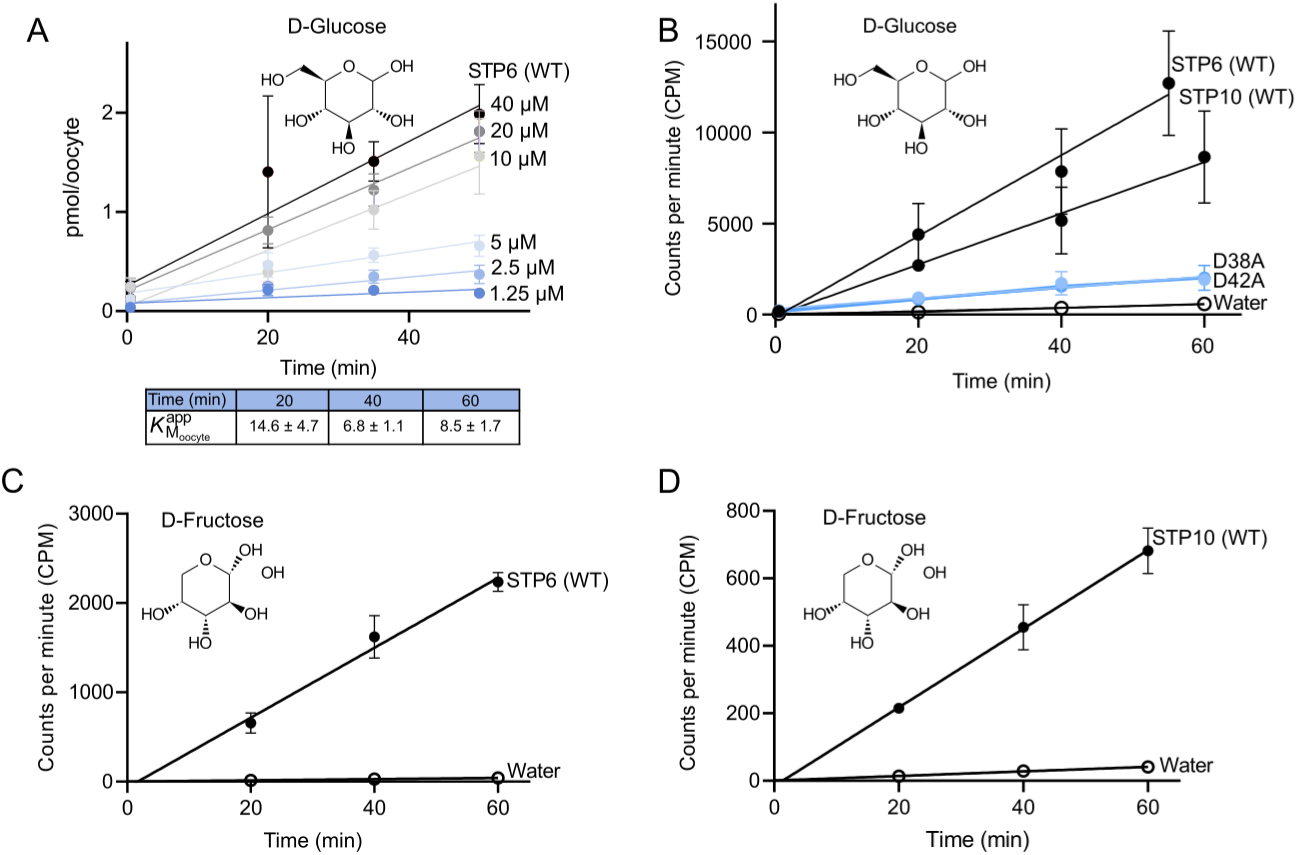
Transport activity tested in *Xenopus oocytes* for STP6 and STP10. **A)** Time dependent uptake of D-glucose by STP6 over a concentration range from 1.25 – 40 µM, used to calculate the apparent KM value shown in figure 3, panel C. The table below shows apparent KM values determined by treating the data as end-point measurements for each of given time points. The variation in *K*_M_ values reflects the observed SD among the individual datapoints used in the analysis shown. **B)** Time dependent uptake of 10 µM D-glucose into *Xenopus oocytes* expressing WT STP10 or WT STP6 (black circles), STP10 D42A or STP6 D38A (blue circles) or water injected oocytes (empty circles). **C)** Time dependent uptake of 10 µM D-fructose into *Xenopus* oocytes expressing WT STP6 (black circles) or water-injected oocytes (empty circles). **D)** Time dependent uptake of 500 µM D-fructose into *Xenopus* oocytes expressing STP10 (black circles) or water injected oocytes (empty circles). All experiments were performed at pH 5 and data points represent mean *±* SD of *n* = 3-5 biologically independent experiments.

**Supplementary Figure 7.**
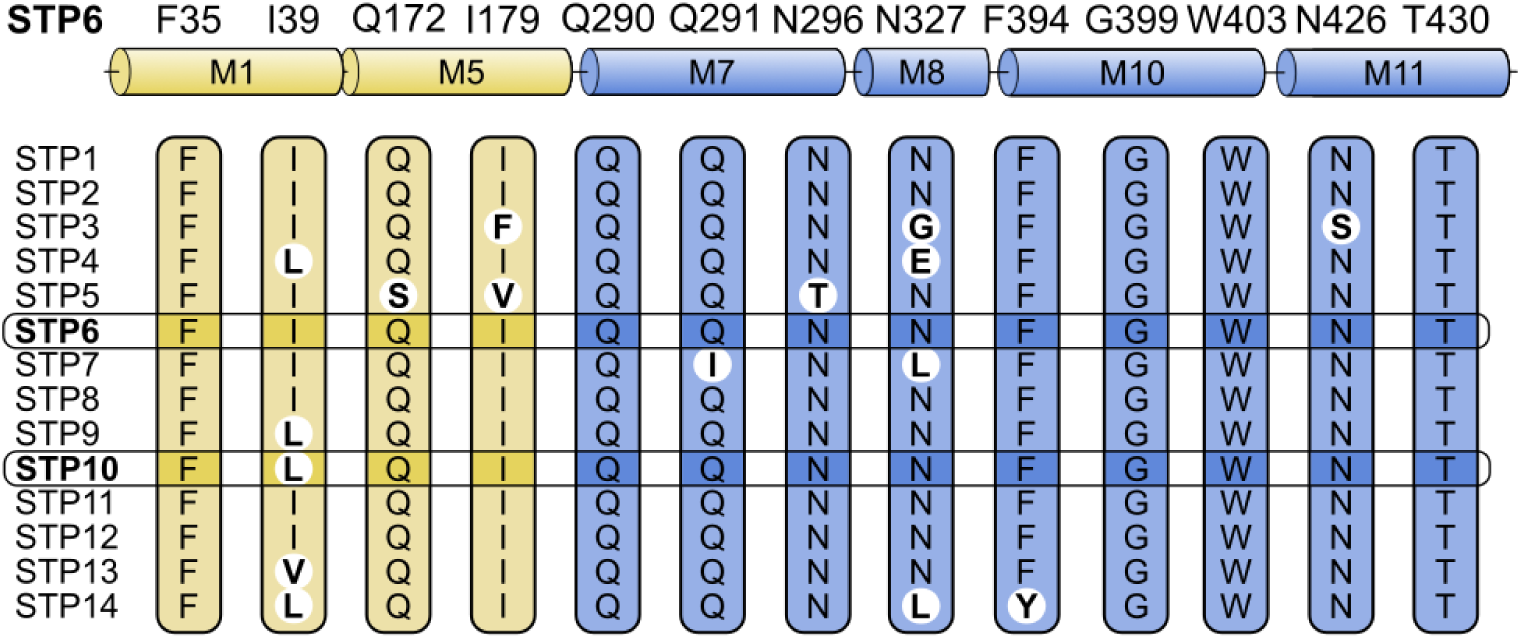
Multiple sequence alignment of residues found in the binding site of *A. thaliana* STPs. Binding site alignment of: STP1 (UniProt accession number P23586), STP2 (UniProt accession number Q9LNV3), STP3 (UniProt accession number Q8L7R8), STP4 (UniProt accession number Q39228), STP5 (UniProt accession number Q93Y91), STP6 (UniProt accession number Q9SFG0), STP7 (UniProt accession number O04249), STP8 (UniProt accession number Q9SBA7), STP9 (UniProt accession number Q9SX48), STP10 (UniProt accession number Q9LT15), STP11 (UniProt accession number Q9FMX3), STP12 (UniProt accession number O65413), STP13 (UniProt accession number Q94AZ2) and STP14 (UniProt accession number Q8GW61). Residues located in the N domain are shown in yellow bars and residues located in the C domain are shown in blue bars. The corresponding helix topology is shown on the top with numbers referring to STP6. Non-conserved residues are highlighted with a white circle and bold font. The STPs concerning this study, STP6 and STP10, are outlined. The only difference in the binding site between STP6 and STP10 is seen on position 39 (STP6) in M5, where an isoleucine instead is a leucine in STP10. STP5 and STP7, which both contain unique sequence deviations, have been reported to be non-functional (7).

**Supplementary Figure 8.**
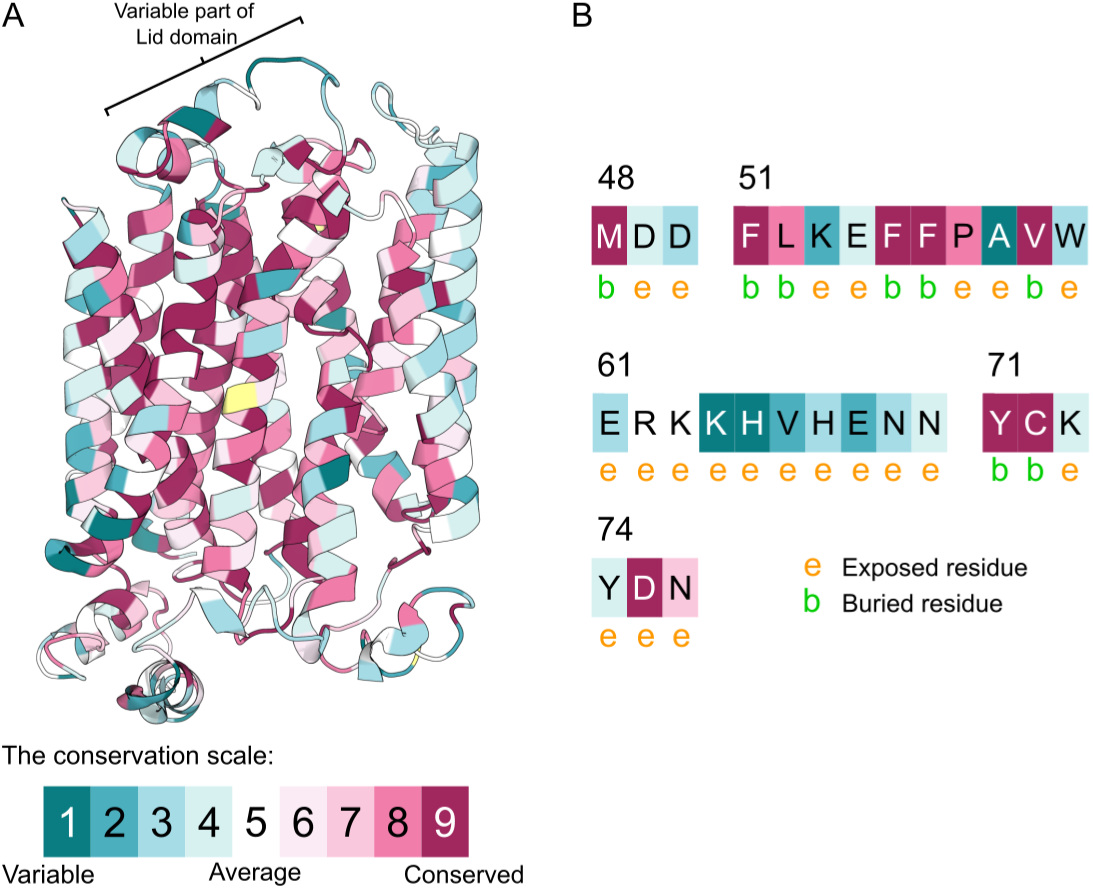
Sequence conservation analysis of *A. thaliana* STPs. **A)** Sequence conservation analysis of the 14 members of the *A. thaliana* STP family represented on the crystal structure of STP6 colored from teal (variable) to bordeaux (conserved). Yellow coloring indicates regions where the data provided was insufficient to calculate a conservation score (region present in < 10% of the data provided). B) Sequence conservation of the STP6 Lid domain along with the predicted exposure of each residue (e = exposed residue, b = buried residue). Exposed residues are seen to be highly variable.

**Supplementary Table 1:** Data collection and refinement statistics.

| Name | STP6 |
| --- | --- |
| State | Inward occluded |
| **Data Collection** |  |
| Space group | P 3<sub>2</sub> 2 1 |
| Cell dimensions |  |
| a, b, c (Å) | 122.22 122.22 139.09 |
| alpha, beta, gamma (deg) | 90 90 120 |
| Monomers per asym. unit. | 1 |
| Wavelength (Å) | 0.9686 |
| Number of reflections measured | 38.166 |
| Number of unique reflections | 19.452 (2813)<sup>a</sup> |
| Resolution (Å) | 84.23-3.20 (3.31-3.20) |
| R<sub>meas</sub> (%) | 14.8 (92.2) |
| Mean I/σ(I) | 7.6 (1.3) |
| CC(1/2) | 98.5 (34.6) |
| Completeness (%) | 95.91 (99.08) |
| Redundancy | 2.0 (1.3) |
| **Refinement** |  |
| Resolution (Å) | 84.23-3.20 (3.31-3.20) |
| No. reflections (work/free) | 19,451 / 922 |
| R<sub>work</sub> (%) | 22.65 |
| R<sub>free</sub> (%) | 25.16 |
| Twin fraction | 0.47 |
| Twin law | -h,-k,l |
| No. of Atoms |  |
| Protein | 3708 |
| Ligands | 117 |
| Average B Factors (Å) |  |
| Overall | 69.1 |
| Protein | 69.1 |
| Ligands | 68.3 |
| RMSD |  |
| Bond lengths (Å) | 0.0047 |
| Bond angles (deg) | 0.87 |
| Validation |  |
| MolProbity score | 1.63 |
| Clashscore | 7.61 |
| Rotamer outliers (%) | 0.00 |
| Ramachandran Plot Statistics |  |
| Favored regions | 96.62 |
| Allowed regions | 3.38 |
| Disallowed regions | 0.00 |
| Deposited model (PDB ID) | 9G11 |

<sup>a</sup> Highest resolution shell is shown in parenthesis.

